# MRBLE-pep measurements reveal accurate binding affinities for B56, a PP2A regulatory subunit

**DOI:** 10.1101/2020.12.16.423088

**Authors:** Jamin B. Hein, Martha S. Cyert, Polly M. Fordyce

## Abstract

Signal transduction pathways rely on dynamic interactions between protein globular domains and short linear motifs (SLiMs). The weak affinities of these interactions are essential to allow fast rewiring of signaling pathways and downstream responses, but pose technical challenges for interaction detection and measurement. We recently developed a technique (MRBLE-pep) that leverages spectrally encoded hydrogel beads to measure binding affinities between a single protein and 48 different peptide sequences in a single small volume. In prior work, we applied it to map the binding specificity landscape between calcineurin and the PxIxIT SLiM (Nguyen et al. 2019). Here, using peptide sequences known to bind the PP2A regulatory subunit B56, we systematically compare affinities measured by MRBLE-pep or isothermal calorimetry (ITC) and confirm that MRBLE-pep accurately quantifies relative affinity over a wide dynamic range while using a fraction of the material required for traditional methods such as ITC.

## Introduction

Cellular signaling transduction pathways are essential for every living organism. At the molecular level, many critical interactions within these pathways involve a globular protein domain binding a short (3-10 amino acid) linear motif (SLiM) within an intrinsically disordered region of a target protein (Neduva and Russell 2006; Dinkel et al. 2016; Davey, Cyert, and Moses 2015). These protein-protein interactions (PPIs) are considered attractive therapeutic targets, and there has been significant recent interest in developing small molecules to specifically target these PPI domains (*e*.*g*. PROTACS) (Paiva and Crews 2019). However, many protein-SLiM interactions remain either uncharacterized or poorly characterized, as they tend to be relatively weak (*K*_d_s in the 1-10 µM range) and are often dynamically regulated via reversible post-translational modifications that modulate binding affinity to alter downstream responses, which complicates their detection, measurement, and characterization. New technologies for discovering, validating, and characterizing protein-SLiM interactions are therefore essential to unlock their therapeutic potential.

High-throughput technologies such as affinity purification coupled to mass spectrometry (Hertz et al. 2016), phage display (Blikstad and Ivarsson 2015; Ueki et al. 2019), and yeast two-hybrid assays (Mu et al. 2014), among others, have dramatically enhanced the pace of candidate protein-SLiM interaction discovery. Furthermore, computational modelling of motif determinants can allow *in silico* screening of entire proteomes to determine motif candidates (Krystkowiak, Manguy, and Davey 2018). However, these high-throughput methods generate large numbers (100s-1000s) of candidate interactors, many of which may be false positives. Validating and quantifying these low affinity binding interactions remains an experimental bottleneck, typically requiring sample- and labor-intensive low-throughput techniques such as isothermal calorimetry (ITC), surface plasmon resonance (SPR), and fluorescence polarization anisotropy, all of which require large amounts of purified protein.

In recent years, several technologies have attempted to bridge this gap between high-throughput, qualitative assays and low-throughput, quantitative assays. Protein microarrays (Meyer et al. 2018) and Luminex bead-based assays (Cook et al. 2019) allow multiplexed measurements of ∼10-1000 interactions in parallel, but do not take place at equilibrium and therefore cannot return thermodynamic binding constants. Microscale thermophoresis and hold-up assay approaches can facilitate such thermodynamic measurements and have successfully been applied towards a variety of systems, including PDZ- and Chromo domain peptide interactions, antibody/antigen interactions, and receptor/ligand interactions (Wienken et al. 2010; Plach, Grasser, and Schubert 2017; Vincentelli et al. 2015). However, they still require fairly high amounts of purified protein.

To complement these approaches, we previously developed MRBLE-pep, a technology that leverages spectrally encoded beads for medium-throughput quantitative measurement of protein-peptide interactions using very small amounts of material (2 µmol in 200 µL for one concentration experiment, or ∼25 µg for B56, the protein analyzed here). To create spectrally encoded beads, we use a microfluidic device to generate porous PEG-DA hydrogel droplets containing specific ratios of 4 different species of lanthanide nanophosphors (LNPs), each of which comprises a unique spectral code (MRBLEs, for <u>M</u>icrospheres with <u>R</u>atiometric <u>B</u>arcode <u>L</u>anthanide <u>E</u>ncoding) (Gerver et al. 2012; Nguyen et al. 2016) Using a custom-built fraction collector (Longwell and Fordyce 2020), we then direct all beads containing a given code to a specified well within a standard multi-well plate **(Fig. 1A)**. After collection, beads are chemically functionalized with Fmoc-glycine and then transferred to a high-throughput peptide synthesizer for Fmoc solid phase peptide synthesis, yielding MRBLE-pep libraries with a unique 1:1 linkage between peptide sequence and embedded spectral code **(Fig. 1A)**. After synthesis, MRBLE-pep bead libraries are pooled and incubated with a fluorescently-labeled protein of interest, followed by imaging to quantify protein binding and identify embedded codes (and thus, the underlying peptide sequence) **(Fig. 1B)**. By repeating MRBLE-pep measurements over multiple protein concentrations, we generate concentration-dependent binding curves, which are fitted to Langmuir isotherms to quantify interaction affinities.

**Figure 1.**
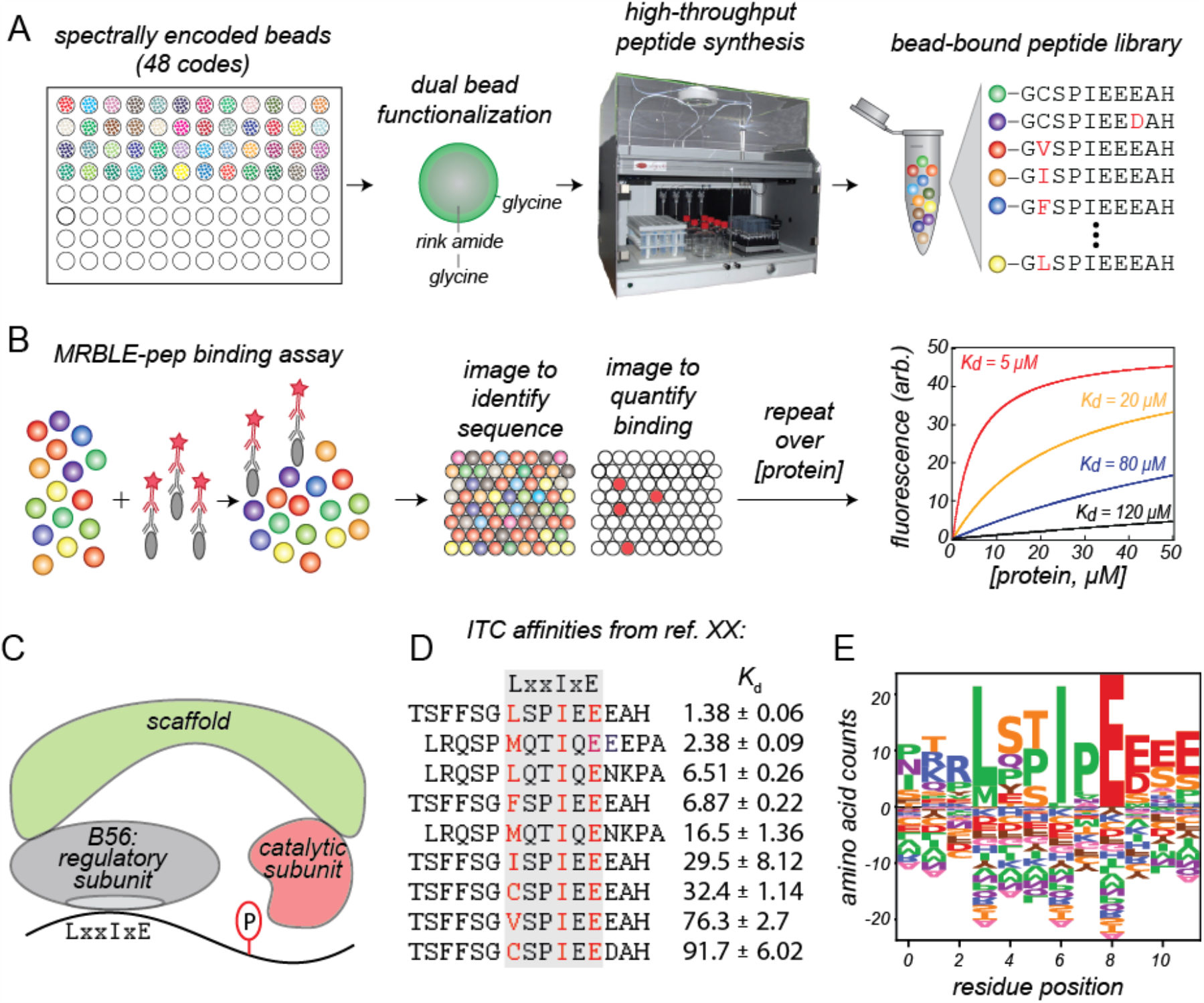
Overview of MRBLE-pep workflow and B56 specificity. **(A)** Schematic showing MRBLE-pep peptide library production. Spectrally encoded hydrogel beads containing unique ratios of lanthanide nanophosphors (LNPs) are functionalized with rink amide/glycine linkers within bead cores and glycine linkers in the outer shell; after functionalization, peptides are synthesized directly on beads with a 1:1 linkage between the peptide sequence identity and embedded spectral code via solid-phase peptide synthesis in a high-throughput peptide synthesizer. **(B)** MRBLE-bound peptide libraries are then pooled and incubated with a protein of interest, primary antibody, and labeled secondary antibody. After binding reactions reach steady state, beads are imaged in the lanthanide channels (to identify the peptide sequence associated with each bead) and in fluorescence channels (to quantify the amount of bead-bound protein). Measurements of binding at multiple concentrations can be globally fit to Langmuir isotherms to quantify interaction affinities. **(C)** Cartoon model of a the heterotrimeric B56-PP2A complex binding a substrate containing the LxxIxE SLiM recognition site. **(D)** Affinities measured by ITC for a set of 9 peptides containing systematic substitutions within the LxxIxE SLiMs from Kif4A and FoxO3 (data from REFERENCE). **(E)** Consensus position-specific specificity matrix generated from known B56 binding sites (data from http://slim.icr.ac.uk/pp2a/index.php?page=instances#28884018) and (Hertz et al. 2016; Wu et al. 2017); logo generated using LogoMaker software (Tareen and Kinney 2020).

In prior work, we applied MRBLE-pep to probe binding interactions between human calcineurin (CN), the Ca2+/calmodulin regulated phosphatase and target of immunosuppressant drugs (Roy and Cyert 2020), and systematic mutations within the known PxIxIT SLiM. CN affinities for six PxIxIT peptide sequences correlated with those previously measured via orthogonal techniques and successfully predicted the strength of signaling inhibition in cells (Nguyen et al. 2019). However, these affinity comparisons were limited, and included several peptides with widely varying affinities reported in the literature. Unfortunately, our own attempts to acquire comparable data for these peptides using established technologies were unsuccessful.

Here, we extend the capabilities of the MRBLE-pep assay and further validate MRBLE-pep-derived affinity measurements using B56, a regulatory subunit of protein phosphatase 2A **(Fig. 1C)** (Ruvolo 2016; McCright and Virshup 1995). In recent work, the Nilsson group and others established that PP2A-B56 substrate specificity relies on a protein-protein interaction between B56 and the LxxIxE SLiM found in various substrates including KIF4A (a kinesin motor protein important for chromosome segregation) and FOXO3 (a transcription factor important for processes such as apoptosis and autophagy) **(Fig. 1D)** (Hertz et al. 2016; Wu et al. 2017). As differences in affinity between B56 and different substrates are thought to dictate the strength and relative timing of downstream signaling responses, the authors quantified the effects of different amino acid substitutions on binding affinities (*K*_d_s) for 8 LxxIxE SLiMs via ITC, revealing a previously unanticipated mode of substrate selection by B56. This set of 50 affinities with a wide dynamic range (spanning from ∼1-100 µM), all measured by a single researcher expert in ITC, provide an ideal comparison set for testing the ability of MRBLE-pep to accurately return affinity information. By directly comparing MRBLE-pep- and ITC-derived affinities for B56 protein interacting with nine distinct peptides, we demonstrate that the MRBLE-pep measurements are in good agreement with ITC literature data. Importantly, the rank order of affinities between different peptides is preserved in MRBLE-pep measurements resulting in the same conclusions in terms of the binding sequence variations while requiring significantly less material.

## Results

To test if MRBLE-pep can return accurate affinity information for an additional PPI system that also requires a different mode of protein detection, we measured binding of purified untagged B56 protein (graciously provided by the Nilsson lab (Hertz et al. 2016)) to two libraries of peptides (containing 22 and 24 peptides, respectively) via MRBLE-pep. To facilitate direct comparison between MRBLE-pep and ITC measurements over a wide dynamic range, each library included 9 peptides with previously measured affinities for B56 spanning 1 to 92 µM **(Fig. 1D)**.

### A generalizable method for fluorescent detection of MRBLE-bound protein

Unlike in the original MRBLE-pep publication (Nguyen et al. 2019) where we detected binding of His-tagged CN to libraries of MRBLE-bound peptides via binding of a fluorescently-labeled anti-His antibody, here we detected binding of untagged B56 protein using a commercially-available anti-B56 primary antibody and an Alexa-647-labeled secondary antibody **(Fig. 2A**,**B)**. Fluorescence images of bead-bound protein revealed a strong bead-associated Alexa-647 signal only in the presence of the B56 protein, establishing that non-specific antibody binding to MRBLE-bound peptides was insignificant **(Fig. 2A**,**B)**. Fluorescent signals for beads coupled to known peptides with high affinity for B56 were >30-fold higher than those for negative controls, establishing that binding was detected with a high dynamic range **(Fig. 2A**,**B)**.

**Figure 2.**
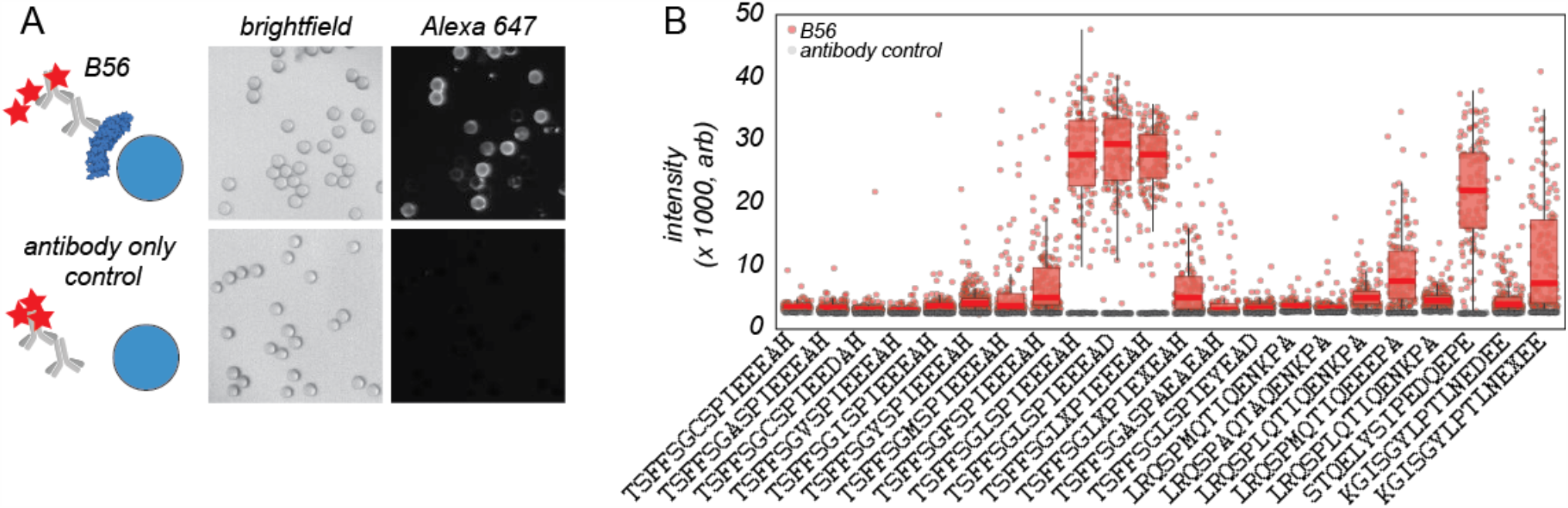
Example images and measured fluorescence intensities for beads incubated with antibodies and B56 or antibodies alone (negative control). **(A)** Schematic of binding assays and example images of beads in the bright field and Alexa 647 fluroescence channels. **(B)** Median binding intensities for each bead (markers) and 95% confidence intervals (box plots) for binding assays with B56 and antibodies (red) or antibodies alone (grey) for a MRBLE-pep assay with 22 different peptide sequences.

### Estimating time-to-equilibrium and verifying appropriate concentrations

Accurately estimating interaction affinities by fitting Langmuir isotherms to observations of concentration-dependent binding relies on two main assumptions. First, interactions must have reached steady state, with no further change in the relative bound and unbound protein fractions over time. Second, the amount of protein ligand available for binding must be in vast excess of the amount of MRBLE-bound peptides in each experiment and significantly higher than expected interaction *K*_d_s (Pollard 2010). For protein-peptide interactions (PPIs) in solution, the time required for a given PPI to reach equilibrium (*K*_eq_) can be loosely approximated by the dissociation rate of the interaction (*k*_off_). For relatively weak interactions (*K*_d_s in the 1-10 µM range), this dissociation rate typically ranges from ∼10^3^-10^5^/s, corresponding to a time to reach equilibrium of ∼0.04 -4 s (Jarmoskaite et al. 2020). In our prior MRBLE-pep experiments, the time required for CN-peptide interactions to reach equilibrium was approximately 14 hours, presumably due to the fact that CN proteins interacting with peptides covalently coupled to the hydrogel matrix were sterically constrained within the porous hydrogel mesh, where the high local concentration of peptide ligands promotes rebinding **(Fig. 2B)**. These slow equilibration times are critical for assay performance, as they make it possible to wash and image beads without loss of weakly bound material during imaging. However, it remained unclear if additional proteins would also require extended times to reach assay equilibrium or whether this property would depend on additional factors such as peptide density, buffer composition, or the identity of the protein itself.

To experimentally test the time required for MRBLE-pep B56 binding to reach equilibrium, we incubated B56 protein at 2 µM with a MRBLE-bound peptide sequence with high affinity for B56 (TSFFSGLSPIEEEAD, code 10) **(Fig. S1-4)**. The reactions required ∼24 hours to reach equilibrium **(Figs. 3C**,**S1-4)**, providing additional support for a model in which protein/peptide dissociation rates are significantly slowed by protein rebinding to other peptides immobilized within the hydrogel matrix. Based on these measurements, we quantified binding for all subsequent B56 MRBLE-pep experiments after 24 hours. To confirm that B56 protein was indeed in vast excess, we first calculated the expected concentrations of B56 protein and available peptides within each binding assay. For a given peptide sequence, we use approximately 100 beads with an estimated peptides density of 10^8^ peptides per bead (Nguyen et al. 2019). We therefore have a peptide concentration of 2.4 x 10^11^ peptides in a 100 μl assay volume for 24 different codes and a protein concentration of 1.86 x 10^13^ at the lowest concentration of protein (31 nmol). For direct experimental validation, we additionally removed the supernatant after 24 hours of incubation and quantified remaining B56 via Western blot, establishing no depletion over the full range of concentrations tested **(Fig. S5)**.

**Figure 3.**
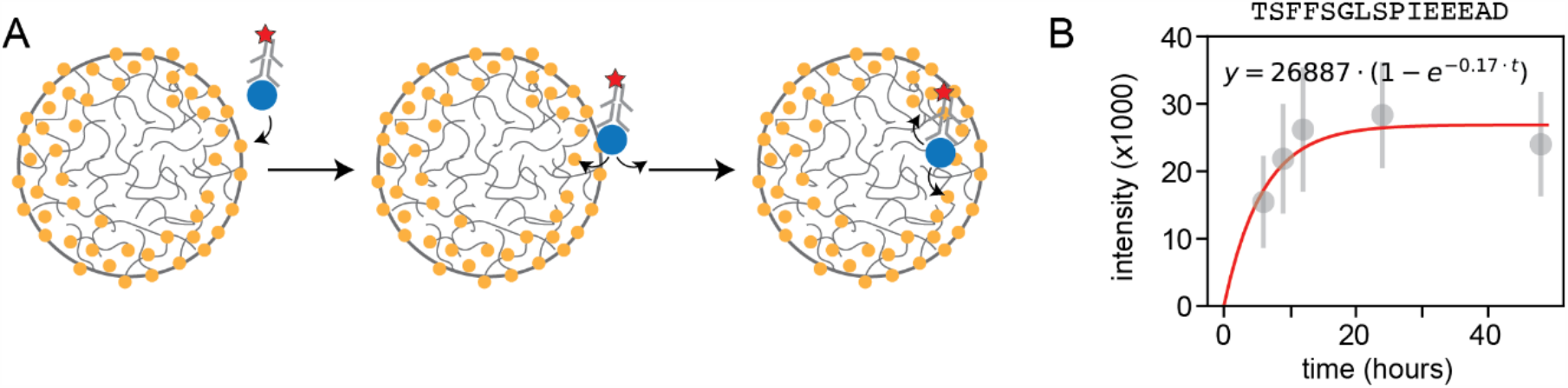
Hydrogel beads reduce dissociation rates to allow equilibrium binding measurements for even weak interactions. **(A)** Cartoon schematic showing proposed mechanism in which proteins binding to bead-bound peptides become trapped in bead hydrogel mesh. **(B)** Time course showing measured intensity as a function of time for B56-antibody complexes interacting with a peptide displayed by MRBLEs hydrogel beads. Markers indicate median intensities, error bars indicate standard deviation, and red line indicates a fit to kinetic binding model (equation and fitted parameters displayed at top of graph).

### Comparing affinities measured across MRBLE-pep technical replicates

Next, for each MRBLE-pep library, we performed 3 technical replicate experiments measuring concentration-dependent binding for B56 protein at 7 concentrations ranging from 31 to 2000 nmol, then globally fit the median intensities for all beads containing a given code at each concentration to a Langmuir isotherm **(Figs. 4A, S6-13)**:

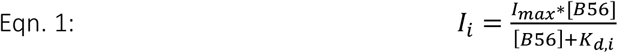

Here, *I*_i_ represents the median fluorescence signal associated with all beads bearing a particular peptide at a given concentration, *I*_max_ is a global saturation value shared across all peptides, [B56] represents the soluble B56 concentration available for binding, and *K*_d,i_ is the binding affinity for a given peptide. By constraining all curves to share a single saturation value for fluorescence intensity, this global fitting procedure returns reliable binding affinity estimates even for peptides with *K*_d_s well above the maximum B56 concentration used in the assay (see **Materials and Methods**). We then convert measured *K*_d_s (absolute affinities) to relative differences in binding energies upon mutation (ΔΔGs) using the following equation:

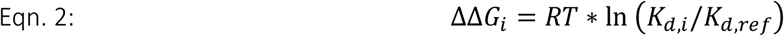

Here, ΔΔG_i_ represents the relative change in binding energy for a given peptide with *K*_d,i_ relative to a ‘reference’ peptide and R and T represent the natural gas constant and temperature, respectively. Pairwise comparisons of measured *K*_d_ and ΔΔG values between replicates for both libraries established that measurements were highly reproducible, with *r*^2^ values of 0.73-0.99 and RMSEs of 0.11-0.37 for *K*_d_s ranging over nearly 3 orders of magnitude and *r*^2^ values of 0.95-0.98 and RMSEs of 0.03-0.65 for ΔΔGs spanning nearly 5 kcal/mol (**Figs. S9**,**S13**). Measurements for the same peptide *across* the two different MRBLE-pep libraries also showed strong agreement (*r*^2^ = 0.86 and RMSE = 0.84 for *K*_d_ comparisons) (**Figs. 4B**). Overall, measured affinities for peptides from library #2 were systematically higher than for library #1, consistent with prior observations that uncertainties in the estimated intensity at saturation can lead to systematic variations in measured affinities. However, relative differences in binding energies remained consistent, particularly when considering systematic mutations within a given peptide backbone (*r*^2^ = 0.83 and RMSE = 0.59 for ΔΔG comparisons, corresponding to <2-fold differences across experiments).

**Figure 4.**
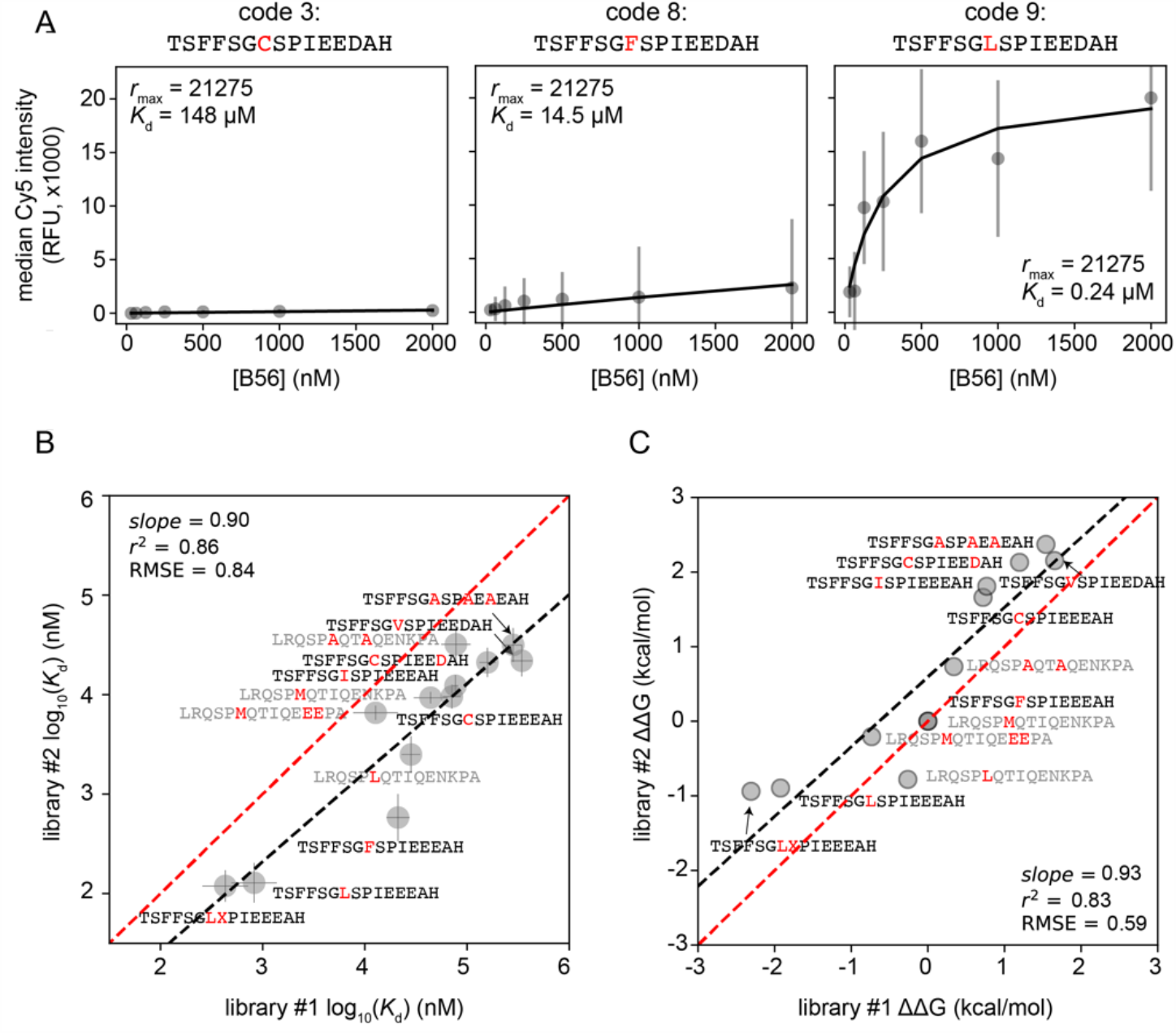
Technical reproducibility of MRBLE-pep replicate binding measurements. **(A)** Concentration-dependent binding data from library #1 for B56 interacting with 3 peptides containing different single amino acid substitutions within the Kif4A LxxIxE SLiM. Grey markers indicate median fluorescence intensities for all MRBLEs displaying a particular peptide sequence at a given concentration; error bars indicate standard deviation of intensities. Solid grey lines denote Langmuir isotherm fits sharing a single maximum intensity saturation value (see **Methods**). **(B)** Measured log_10_-transformed binding affinities for 12 peptides measured via 2 independent full technical MRBLE-pep replicate experiments. Grey markers indicate the mean of 3 independent MRBLE-pep replicates for a given library; error bars denote the associated standard deviation. The red dashed line indicates a 1:1 relationship and the black dashed line indicates the results of a linear regression to the log_10_-transformed *K*_d_ values. **(C)** Measured ΔΔGs for 12 peptides measured across 2 full technical MRBLE-pep replicates. Grey markers indicate the mean of 3 independent MRBLE-pep replicates for a given library; error bars denote the associated standard deviation. The red dashed line indicates a 1:1 relationship and the black dashed line indicates the results of a linear regression.

### Comparing affinities measured via MRBLE-pep and ITC

Next, we sought to directly compare affinities measured via MRBLE-pep and ITC (isothermal calorimetry). While ITC has traditionally been considered the ‘gold standard’ method for measuring *K*_d_s for protein interactions, these experiments require large amounts of highly purified protein, particularly for weak protein-peptide interactions (900 μg per protein/peptide interaction for *K*_d_s in the low micromolar range). MRBLE-pep measurements require significantly less material (25 μg for 22 protein/peptide interaction measurements, or 1 μg per interaction) and labor (as multiple protein/peptide interactions can be assessed in a single experiment), but to date have only been shown to return thermodynamic binding information for a single protein (CN).

Measured affinities (*K*_d_s) for B56 binding to 12 peptides containing systematic substitutions within the LxxIxE motif found within Kif4A or FoxO3 showed strong correlation between the two methods (*r*^2^ = 0.83 and 0.70 for libraries 1 and 2, respectively) (**Fig. 5A**). Measured affinities from library #1 showed better agreement with ITC-derived affinities (RMSE = 0.37 and 0.61 for libraries 1 and 2, respectively), largely due to a single measurement for a FoxO3 variant containing additional negatively charged residues at the C-terminal end of the sequence (LRQSPMTIQEEEPA) (**Fig. 5B, Table S1**). Overall, library 2 measurements returned systematically higher affinities, potentially due to small variations in the surface peptide density of displayed peptides between replicates. However, measurements of relative changes in binding affinities upon mutation (ΔΔGs) showed strong correlation for both libraries (*r*^2^ = 0.93 and 0.94 for both libraries, with RMSEs of 0.48 and 0.31, respectively, and the rank order of preferred sequences remained largely unchanged. Together, these comparisons establish that MRBLE-pep returns binding affinities with similar accuracy to ITC but using 1/500th of the material.

**Figure 5.**
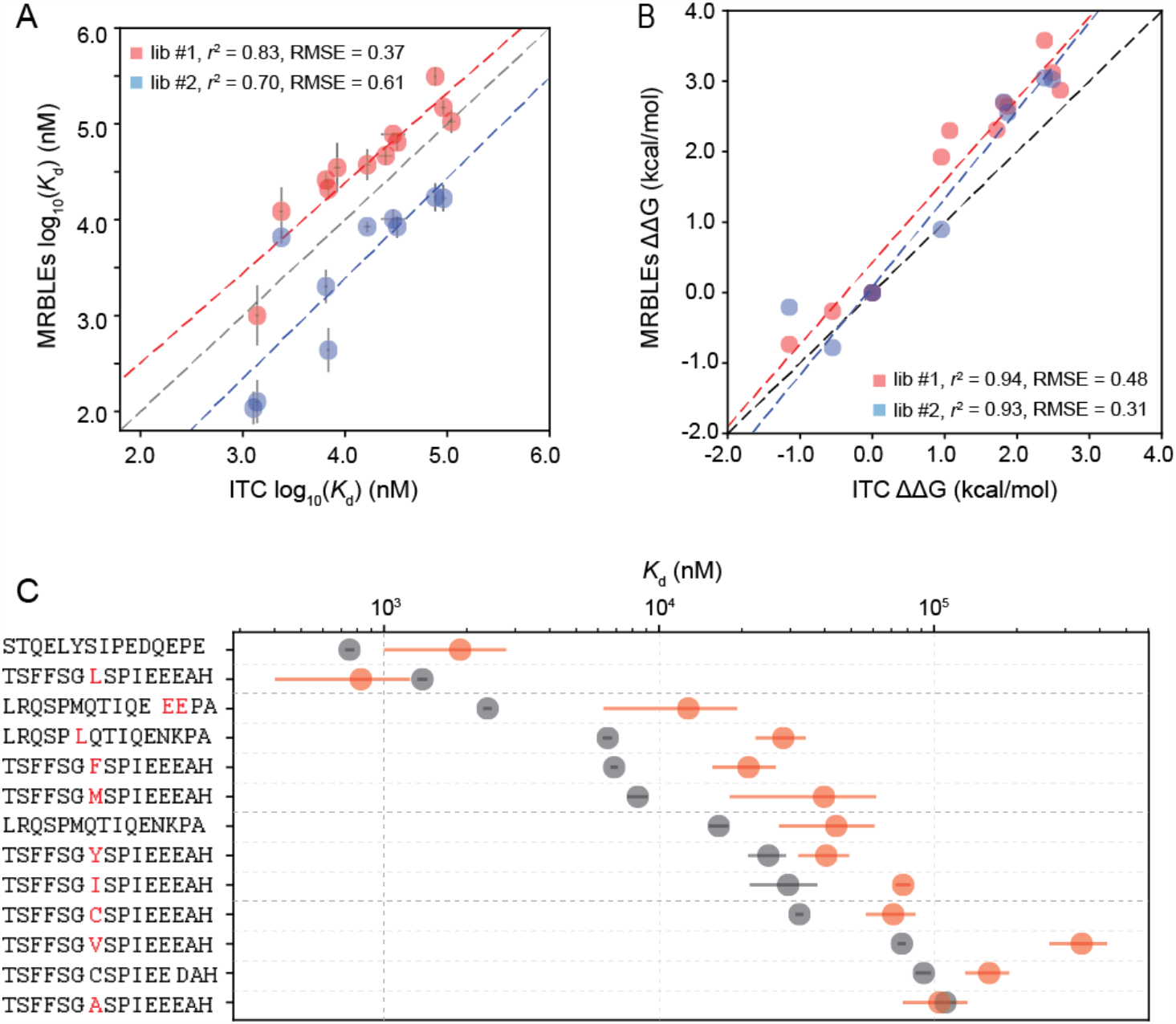
Comparison between MRBLE-pep and ITC affinity measurements. **(A)** Comparison of absolute affinities (*K*_d_s) measured via MRBLE-pep libraries #1 (red) and #2 (blue) and ITC (data from Hertz*,Kruse*, *et al*.) for 9 peptides derived from Kif4A and FoxO3 SLiM recognition motifs present in all 3 libraries (see **Fig. 1D**). Each MRBLE-pep measurement represents the median of 3 technical replicates and error bars indicate the standard error of the mean; for ITC measurements, error bars indicate the uncertainties returned from Langmuir isotherm fits. (**B**) Comparison of relative differences in affinity (ΔΔGs) measured via MRBLE-pep libraries #1 (red) and #2 (blue) and ITC (data from Hertz*,Kruse*, *et al*.) for 9 peptides derived from Kif4A and FoxO3 SLiM recognition motifs present in all 3 libraries (see **Fig. 1D**). (**C**) Comparison between affinities measured by MRBLE-pep library #1 (red) and ITC (data from (Hertz et al. 2016)) for 13 peptides present in both libraries.

### Applying MRBLE-pep to investigate how mutations within LxxIxE SLiMs affect B56 binding

Having established the accuracy of MRBLE-pep, we applied it to investigate the effects of systematic substitutions within the LxxIxE SLiM motifs present within Kif4A, FoxO3, GEF-H1, and Rac-GAP1. In particular, we investigated the influence of different amino acids in the first position and those surrounding the acidic last position of the motif. In addition, we probed how phosphorylation at different positions in close proximity to key positions affected binding affinities. (**Fig. 6**). Within Kif4A, the LxxIxE motif appears as a CxxIxE sequence (**Fig. 6A**). As expected, making a C to L substitution at the first position within the SLiM (which alters the site to look more like the known consensus) enhances affinity by ∼2.5 kcal/mol (**Fig. 6A**). Overall, the L residue is strongly preferred at this position, with F and M substitutions also moderately enhancing binding relative to the native C. In the presence of mutations to this first position, mutating the additional I and E conserved residues reduces binding to background levels. Surprisingly, substituting the L or the E position with a Y in a high affinity LxxIxE motif did not completely abolish binding (reduction of ∼2 or ∼3 kcal/mol, respectively), suggesting that SLiMs containing an L in this first position or multiple acidic residues at the C-terminal boundary may be able to tolerate substitutions at other conserved positions while maintaining biological function (**Fig 6A**). However, future *in vivo* work is required to test this hypothesis.

**Figure 6.**
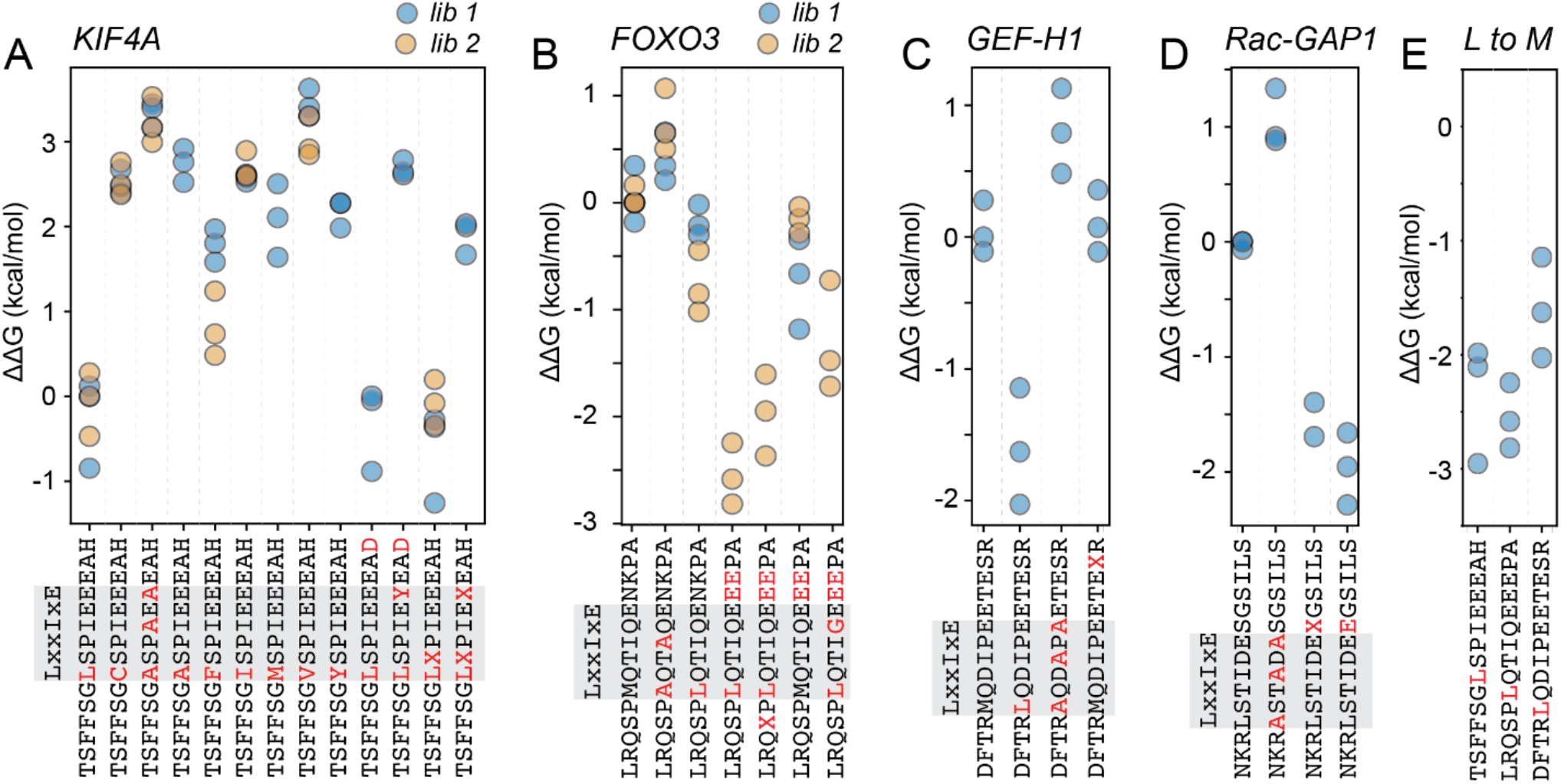
MRBLE-pep measurements quantifying the effects of systematic amino acid substitutions within the Kif4A, FoxO3, GEF-H1, and Rac-GAP1 LxxIxE SLiM on B56 binding affinity. **(A)** Measured ΔΔGs for systematic substitutions within the Kif4A LxxIxE SLiM relative to the median TSFFSGLSPIEEEAH value. **(B)** Measured ΔΔGs for systematic substitutions within the FoxO3 LxxIxE SLiM relative to the median WT (LRQSPMQTIQENKPA) value. **(C)** Measured ΔΔGs for systematic substitutions within the GEF-H1 LxxIxE SLiM relative to the median WT (LDFTRMQDIPEETESR) value. **(D)** Measured ΔΔGs for systematic substitutions within the Rac-GAP1 (NKRLSTIDESGSILS) value. **(E)** Measured ΔΔGs for M to L substitutions within 3 different peptide backbones (Kif4A, FoxO3, and GEF-H1).

In recent work, the Nilsson lab also reported that phosphorylation generally increases B56 binding affinities when adjacent to either the glutamic acid residues at the C-terminus of the LxxIxE SLiM or to the conserved N-terminal leucine. Here, we find that the effects of phosphorylation are strongly context-dependent: while a single phosphorylated residue adjacent to the conserved leucine leads to slightly enhanced affinities, the addition of a second phosphorylated residue at the C-terminus of the Kif4A SLiM dramatically decreases binding (**Fig. 6A**). By contrast, the addition of a phosphorylated serine N-terminal to the FoxO3 LxxIxE motif reduces binding (**Fig. 6B**), phosphorylation of residues adjacent to the glutamic acids within the GEF-H1 SLiM has no effect on affinities, and phosphorylation immediately C-terminal to the Rac-GAP1 LxxIxE SLiM dramatically enhances binding (**Fig. 6C**,**D**). Interestingly, an additional E after the obligatory E seems to increase affinity more than a phosphorylated serine (**Fig. 6D**). Furthermore, we observed a strong decrease in binding affinity when mutating position five (LxxI<u>x</u>E) from D to G in FoxO3, indicating that this position could be important for fine tuning the affinity between B56 and its binding partners (**Fig. 6B**); future experiments are required to systematically probe how substitutions at this position alter affinity. Together, these results highlight the importance of local sequence context in determining binding strength, an observation further supported by the fact that M to L substitutions within a variety of different peptide backbones have different effects (**Fig. 6E**).

## Discussion

In prior work, we described the development of the MRBLE-pep assay and demonstrated that MRBLE-pep was able to quantitatively profile the binding specificity landscape for CN interacting with a library of ∼400 peptides containing systematic substitutions within the PxIxIT SLiM. Here, we further validate that MRBLE-pep agrees with previous ITC measurements in reporting the relative effects of amino acid substitutions on binding affinities of an additional SLiM (LxxIxE) for its receptor (B56), while using a fraction of the required material and requiring a fraction of the time. Moreover, we establish that protein binding can be accurately quantified using an unlabeled primary antibody and fluorescently-labeled secondary antibody, expanding the capabilities of the assay to examine proteins that are difficult to tag and/or purify.

While technical replicates using independent MRBLE-pep libraries showed systematic shifts in raw affinities (**Fig. 5**), we establish that measured ΔΔGs and the rank order of peptide sequence preferences are highly reproducible. We speculate that these systematic shifts may result from differences in the density of displayed peptides after on-bead synthesis. In recent work, we described a dramatically simplified method for encoded bead production and functionalization (MRBLEs 2.0) (Feng et al. 2020), and we anticipate that peptide synthesis on beads produced via this new pipeline will yield more consistent density coverage. Furthermore, by simplifying the technology we hope that these improvements will allow researchers to more broadly adopt MRBLE-pep to systematically characterize SLiM binding determinants and to validate candidate interactions identified by high-throughput phage-display, yeast-two-hybrid or *in silico* screens.

## Materials and Methods

### MRBLE production and peptide synthesis

We produced MRBLEs spectrally encoded hydrogel beads as described previously (Hein et al. 2020; Nguyen et al. 2019).

### B56 expression and purification

B56 was a kind gift from the lab of Jakob Nilsson and was expressed and purified as described previously (Hertz et al. 2016).

### MRBLE-pep time-to-equilibrium assays

To reduce nonspecific binding, we ‘blocked’ beads by incubating approximately 500 beads per code with PBST (0.1 % Tween-20) and 5 % BSA for 1 hour at RT. To allow labeled antibody to bind B56 protein prior to bead assays, we preincubated 2 µmol recombinant B56 protein with 1 µmol mouse anti-B56 antibody (Clone 23/B56α BD biosciences, 610615) and 1 µmol goat anti-mouse IgG Alexa Fluor 647 (Thermo Fisher A-21235) in 500 µL binding buffer (50 mM Tris pH = 7.5, 150 mM NaCl, 0.1% TWEEN 20) for one hour at 4 °C. Prior to beginning binding assays, we washed beads once with binding buffer and divided them into five 200 µl Eppendorf tubes. After decanting the binding buffer, we added 100 µL of the B56 protein and anti-B56 and anti-mouse IgG antibody complex. At specified times, we removed protein-antibody complexes, washed beads once with 100 µL binding buffer, and resuspended in 20 µL PBST prior to imaging.

### Western blotting to test for depletion of B56 after binding assays

Each gel electrophoresis reaction used 6 µL of unbound B56 protein-antibody complex and 2 µL of 4x SDS-PAGE sample buffer (Thermo Fisher). After heating the samples to 95 C for 10 minutes, we loaded samples on a SDS-PAGE gel (4–20% Precast Protein Gels, Bio-Rad #4561095DC) and separated the protein at 100 V for 60 min. We then blotted the separated proteins onto a PVDF membrane for 90 min at 400 mA using a Bio-Rad wet blotting system. The membrane was blocked with 3 % BSA in TBS-T for 30 min at room temperature with gentle agitation. We removed the buffer and added fresh TBS-T with 3% BSA and Goat anti-Mouse IR-Dye 700 (1:5000 diluted) to obtain a background signal from the Mouse anti-B56 antibody present in the samples. After 2 hours incubation with gentle agitation at room temperature, we washed the blot three times for five minutes with TBS-T, dried, and scanned using the Odyssey CLx imaging system (Licor Biosciences). The membrane was incubated again with 3 % BSA in TBS-T for 30 min at room temperature. The buffer was exchanged and 500 ng of Mouse anti-B56 antibody was added and incubated for 2 hours at room temperature. The membrane was again washed 3 times for 5 minutes with TBS-T and then we added fresh buffer containing 1:5000 diluted Goat anti-Mouse IR-Dye 700 and incubated for 1 hour at room temperature. After additional washing, the membrane was scanned using the Odyssey CLx imaging system.

### MRBLE-pep concentration-dependent binding assays

Each concentration-dependent binding assay used approximately 700 beads per code. To begin the assay, we blocked beads with PBST (0.1 % Tween-20) and 5 % BSA for 1 hour at RT to reduce non-specific binding. During this blocking step, we also preincubated 200 µL of 2 µmol recombinant B56 protein with 1 µmol mouse anti-B56 antibody (Clone 23/B56α BD biosciences, 610615) and 1 µmol goat anti-mouse IgG Alexa Fluor 647 (Thermo Fisher A-21235) for one hour at 4 degrees to form B56/antibody complexes. Next, we washed beads once with 700 µl binding buffer and then divided them into seven 200 µl Eppendorf tubes to yield approximately 100 beads per code per assay. B56/antibody complexes were diluted from 2 µmol to the appropriate assay concentration using binding buffer containing 10 % glycerol and added to Eppendorf tubes containing beads to a final reaction volume of 100 µL. After 24 hours, we removed the unbound protein-antibody complexes, washed beads once with 100 µL of binding buffer, and resuspended in 20 µL PBST.

### MRBLE-pep imaging

We imaged all beads exactly as previously described (Nguyen et al. 2019; Hein et al. 2020; Harink et al. 2019). At the end of this analysis, we quantified a median intensity for all pixels associated with this bead annulus for each bead [*source data*].

### Global fitting of concentration-dependent binding curves

To generate binding curves for each code, we first plotted the median of all median bead annulus fluorescence intensities as a function of soluble B56 concentration. To determine the *K*_d_ values for each protein/peptide interaction, we then globally fit the data for all codes to a single-site binding model(Maerkl and Quake 2007; Fordyce et al. 2010; Aditham et al. 2020) :

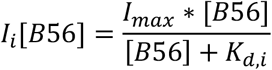

Here, *I*i[B56] denotes the measured bead-associated Cy5 intensity as a function of soluble B56 concentration for a given code, *I*max is a global constant shared across *all* codes corresponding to the maximum bead-associated Cy5 intensity at saturating [B56], [B56] denotes the concentration of soluble B56 available for binding, and *K*d,i denotes the dissociation constant (*K*d) for a given B56-peptide complex. This global fitting procedure assumes that: (1) the binding interaction is measured at steady-state, and (2) the stoichometry of binding remains constant across all B56-peptide interactions (making it possible to estimate *K*_d_s even for very weak interactions with *K*_*d*_ values greater than the maximum soluble B56 concentration probed in the assay.

To calculate ΔΔGs across a given experiment, we used the following formula:

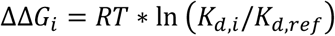

where R is the gas constant (1.987 · 10-3 kcal/(K · mol)), T is 298 K, and *K*_d,i_ is the median of the returned *K*_d_s from technical replicates of concentration-depending binding experiments for a protein-peptide interaction of interest and *K*_d,ref_ is the median of the returned *K*_d_s from technical replicates of concentration-depending binding experiments for a reference protein-peptide interaction.

### Comparing MRBLE-pep technical replicates for a given library

To compare measurements between technical replicates using a given MRBLE-pep library, we performed pairwise comparisons between each replicate and evaluated concordance via a linear regression of: (1) log_10_-transformed *K*_d_ values for each peptide returned from the global fitting procedure described above, and (2) calculated ΔΔG values relative to a single reference peptide.

### Comparing MRBLE-pep measurements across libraries

To compare measurements across MRBLE-pep libraries, we directly compared: (1) the median log_10_-transformed *K*_d_ value from the 3 technical replicates of each library and (2) ΔΔG values calculated from these median *K*_d_ values. In each case, we again evaluated agreement via a linear regression.

### Comparing MRBLE-pep measurements with ITC

To compare measurements between MRBLE-pep and ITC, we again directly compared the median log_10_-transformed *K*_d_ value from the 3 technical replicates of each library with prior measurements from the Nilsson lab (Hertz et al. 2016).

## Acknowledgements

P.M.F. and M.S.C. gratefully acknowledge the support of a Stanford Bio-X Interdisciplinary Initiatives Seed Grant. This work has been supported by NIH grant 1DP2GM123641 awarded to P.M.F., and P.M.F. is also a Chan Zuckerberg Biohub Investigator. JBH was funded by grant NNF17OC0025404 from the Novo Nordisk Foundation and the Stanford Bio-X Program. M.S.C. is funded by NIH grant R35GM136243. P.M.F. previously co-authored a patent describing production of spectrally encoded beads via ratiometric barcode lanthanide encoding (WO 2014/031902 A2).

