## Supplementary Information for "MRBLE-pep measurements reveal accurate binding affinities for B56, a PP2A regulatory subunit"

**Table S1**: Mean $K_d$ (nM) and mean ΔΔG (kcal/mol) for peptides calculated from triplicate measurements for both libraries.

| Code | Sequence | Mean (Kd) | SE (Kd) | Mean (ΔΔG) | SE (ΔΔG) | Library |
|---|---|---|---|---|---|---|
| 1 | TSFFSGCSPIEEEAH | 71073.44 | 10306.54 | 2.82 | 0.23 | 1 |
| 2 | TSFFSGASPIEEEAH | 104654.44 | 19522.14 | 3.04 | 0.19 | 1 |
| 3 | TSFFSGCSPIEEDAH | 158673.79 | 20642.92 | 3.30 | 0.28 | 1 |
| 4 | TSFFSGVSPIEEEAH | 344022.43 | 57937.01 | 3.76 | 0.28 | 1 |
| 5 | TSFFSGISPIEEEAH | 77163.64 | 3446.67 | 2.88 | 0.31 | 1 |
| 6 | TSFFSGYSPIEEEAH | 40576.78 | 6039.53 | 2.49 | 0.19 | 1 |
| 7 | TSFFSGMSPIEEEAH | 39828.93 | 15393.48 | 2.39 | 0.12 | 1 |
| 8 | TSFFSGFSPIEEEAH | 21125.87 | 3881.26 | 2.10 | 0.19 | 1 |
| 9 | TSFFSGLSPIEEEAH | 824.49 | 299.18 | 0.07 | 0.02 | 1 |
| 10 | TSFFSGLSPIEEEAD | 719.33 | 248.05 | 0.00 | 0.00 | 1 |
| 11 | TSFFSGLXPIEEEAH | 430.93 | 157.18 | -0.32 | 0.03 | 1 |
| 12 | TSFFSGLXPIEXEAH | 25598.31 | 4459.53 | 2.21 | 0.17 | 1 |
| 13 | TSFFSGASPAEAEAH | 283395.36 | 38820.16 | 3.64 | 0.35 | 1 |
| 14 | TSFFSGLSPIEYEAD | 92865.40 | 8706.23 | 2.99 | 0.25 | 1 |
| 15 | LRQSPMQTIQENKPA | 44031.44 | 11871.24 | 2.51 | 0.29 | 1 |
| 16 | LRQSPAQTAQENKPA | 77891.82 | 18278.86 | 2.86 | 0.28 | 1 |
| 17 | LRQSPLQTIQENKPA | 27692.73 | 3245.71 | 2.27 | 0.29 | 1 |
| 18 | LRQSPMQTIQEEEPA | 12766.19 | 4588.22 | 1.73 | 0.12 | 1 |
| 19 | LRQSPLQTIQENKPA | 27809.86 | 4479.58 | 2.27 | 0.29 | 1 |
| 20 | STQELYSIPEDQEPE | 1893.03 | 632.36 | 0.59 | 0.06 | 1 |
| 21 | KGISGYLPTLNEDEE | 43087.35 | 10136.94 | 2.50 | 0.16 | 1 |
| 22 | KGISGYLPTLNEXEE | 13148.79 | 4776.67 | 1.71 | 0.05 | 1 |
| 1 | DFTRLQDIPEETESR | 755.78 | 313.65 | 0.43 | 0.04 | 2 |
| 2 | DFTRMQDIPEETEXR | 11812.68 | 2771.72 | 2.14 | 0.09 | 2 |
| 3 | DFTRAQDAPAETESR | 39734.00 | 12185.37 | 2.83 | 0.21 | 2 |
| 4 | DFTRMQDIPEETESR | 10706.87 | 2184.70 | 2.09 | 0.12 | 2 |
| 5 | NKRLSTIDEXGSILS | 311.00 | 76.28 | 0.18 | 0.06 | 2 |
| 6 | NKRLSTIDEEGSILS | 161.85 | 47.09 | -0.43 | 0.04 | 2 |
| 7 | NKRASTADASGSILS | 25288.43 | 6720.82 | 2.58 | 0.12 | 2 |
| 8 | NKRLSTIDESGSILS | 3947.35 | 139.40 | 1.52 | 0.20 | 2 |
| 9 | LRQSPLQTIGEEEPA | 1216.59 | 635.59 | 0.67 | 0.12 | 2 |
| 10 | LRQXPLQTIQEEEPA | 346.25 | 119.03 | 0.00 | 0.00 | 2 |
| 11 | LRQSPLKTIKEEEPA | 54.63 | 18.22 | -1.08 | 0.08 | 2 |
| 12 | LRQSPLQTIQEEEPA | 124.51 | 35.18 | -0.58 | 0.06 | 2 |
| 13 | LRQSPMQTIQEEEPA | 6576.38 | 773.96 | 1.82 | 0.15 | 2 |
| 14 | LRQSPLQTIQENKPA | 2501.93 | 753.05 | 1.20 | 0.08 | 2 |
| 15 | LRQSPAQTAQENKPA | 32042.00 | 9690.55 | 2.71 | 0.16 | 2 |
| 16 | LRQSPMQTIQENKPA | 9325.89 | 886.41 | 2.03 | 0.18 | 2 |
| 17 | TSFFSGASPAEAEAH | 31616.29 | 8755.70 | 2.71 | 0.18 | 2 |
| 18 | TSFFSGLXPIEEEAH | 119.18 | 30.79 | -0.60 | 0.07 | 2 |
| 19 | TSFFSGLSPIEEEAH | 128.32 | 41.73 | -0.58 | 0.02 | 2 |
| 20 | TSFFSGFSPIEEEAH | 581.48 | 224.80 | 0.30 | 0.06 | 2 |
| 21 | TSFFSGISPIEEEAH | 12282.41 | 2181.89 | 2.18 | 0.15 | 2 |
| 22 | TSFFSGVSPIEEEAH | 21912.73 | 5620.81 | 2.50 | 0.11 | 2 |
| 23 | TSFFSGCSPIEEDAH | 21088.21 | 5071.72 | 2.48 | 0.12 | 2 |
| 24 | TSFFSGCSPIEEEAH | 9551.44 | 1878.54 | 2.02 | 0.12 | 2 |

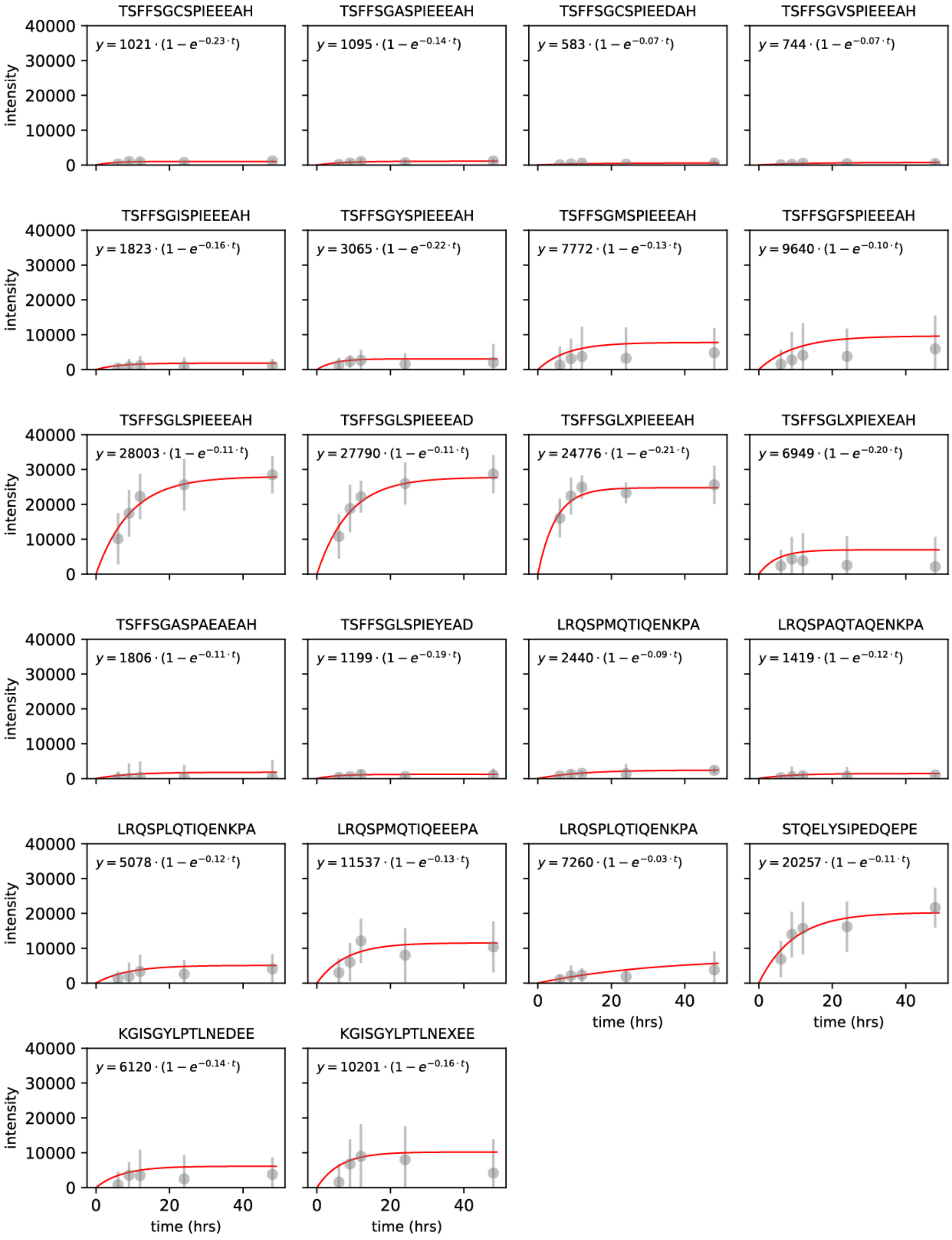


**Fig. S1**. Measurement of time required to reach binding equilibrium for all peptides in MRBLE-pep library 1. Grey markers indicate median bead intensity at a given time point, error bars indicate the standard deviation, and the red line indicates a fit to a kinetic binding curve (equation and fit parameters given at top).

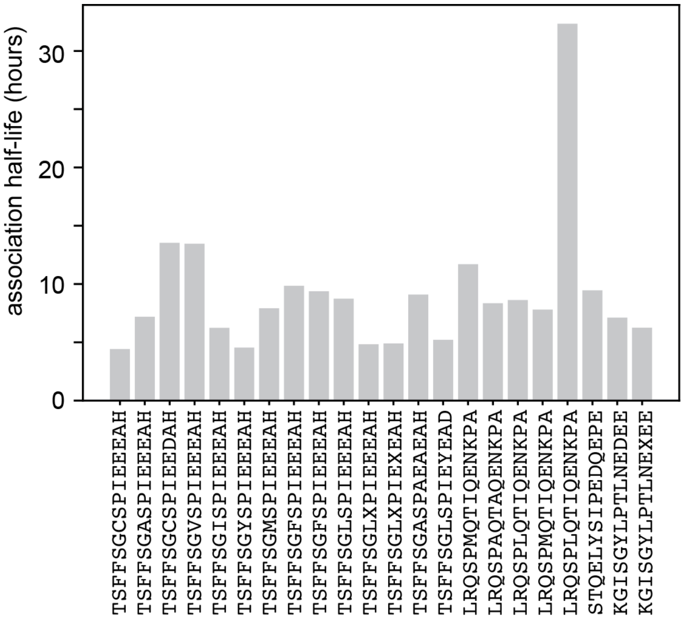


**Fig. S2**. Association half-life for all peptides from MRBLE-pep library 1 (determined from kinetic binding fit parameters).

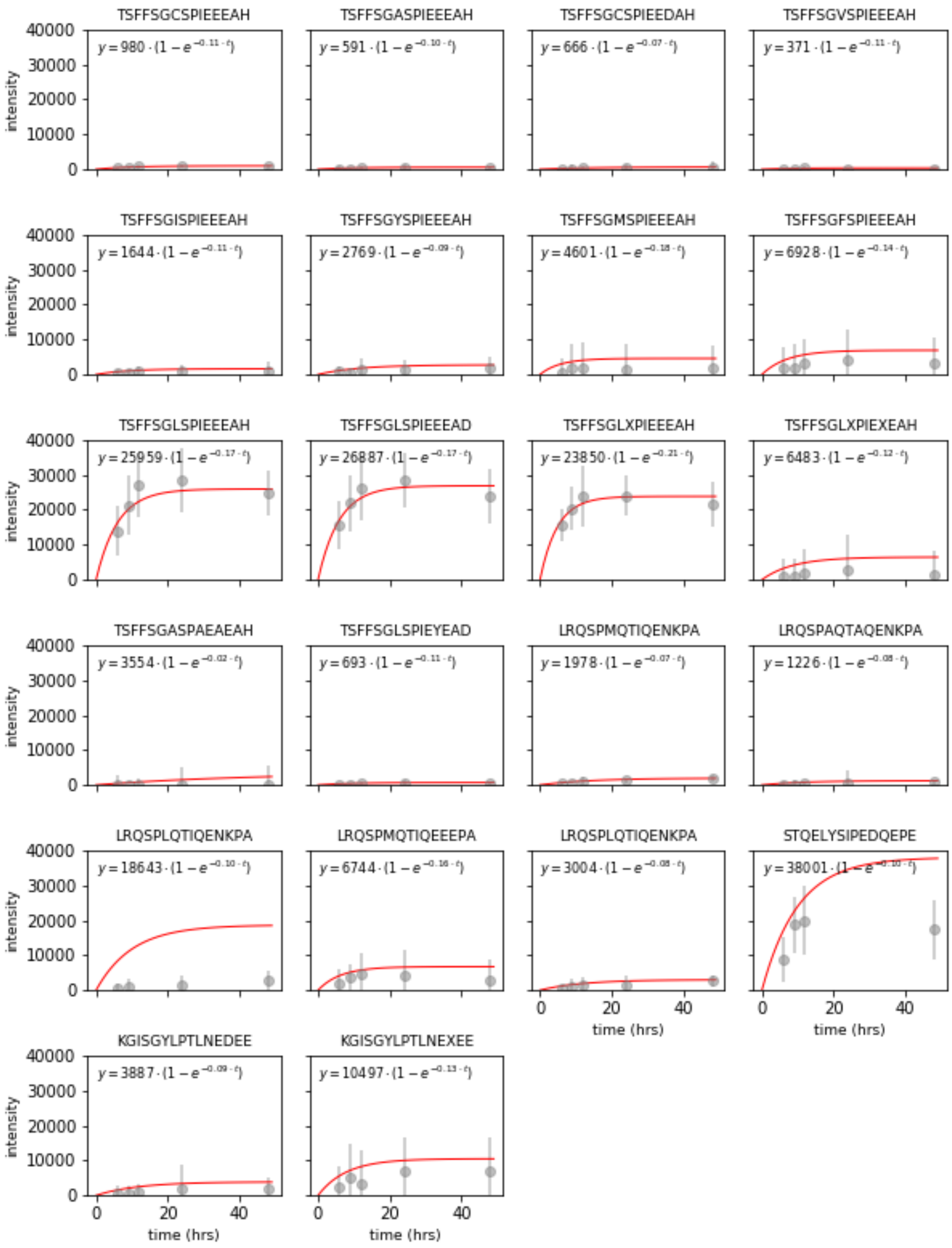


**Fig. S3**. Measurement of time required to reach binding equilibrium for all peptides in MRBLE-pep library 2. Grey markers indicate median bead intensity at a given time point, error bars indicate the standard deviation, and the red line indicates a fit to a kinetic binding curve (equation and fit parameters given at top).

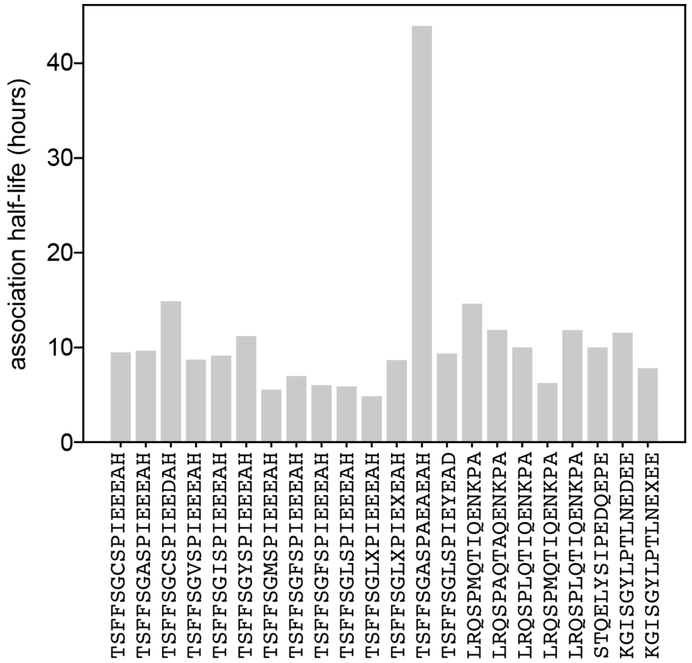


**Fig. S4**. Association half-life for all peptides from MRBLE-pep library 2 (determined from kinetic binding fit parameters).

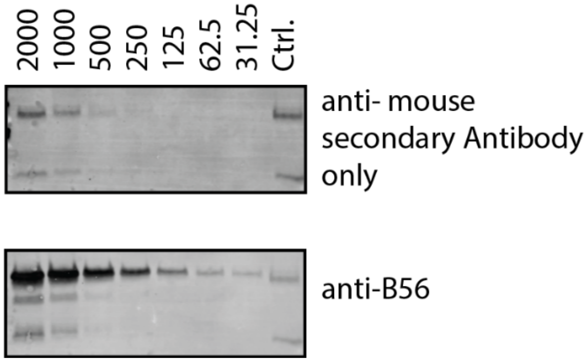


**Fig. S5**. Western Blot with the unbound fractions of a MRBLE-pep concentration experiment. The unbound fraction from each concentration of a MRBLE-pep concentration experiment was probed for unbound B56 protein.

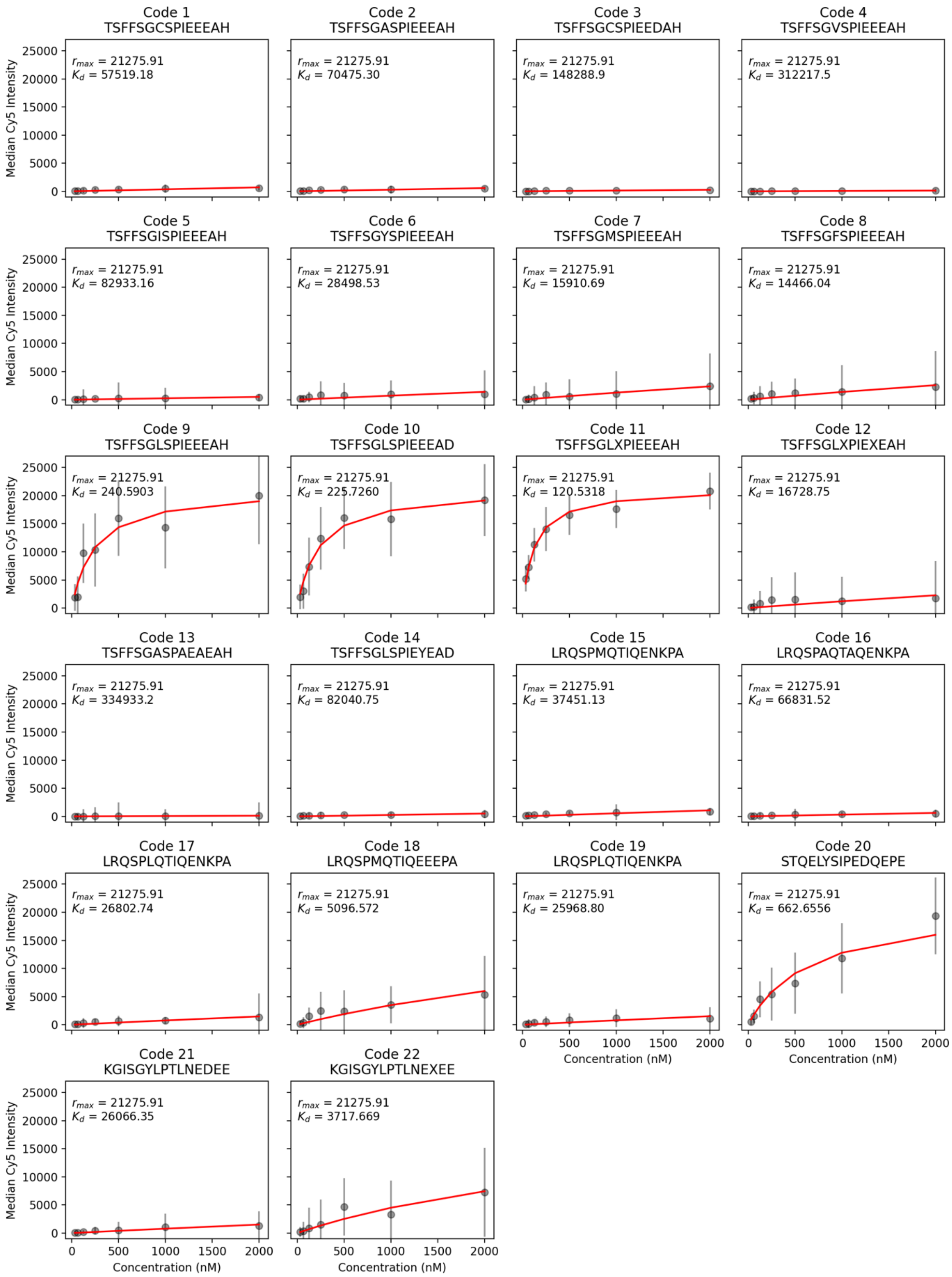


**Fig. S6**. Concentration-dependent binding data (grey markers, median intensity and standard deviation over all beads) and associated Langmuir isotherm fits (red lines) for MRBLE-pep library 1, replicate #1.

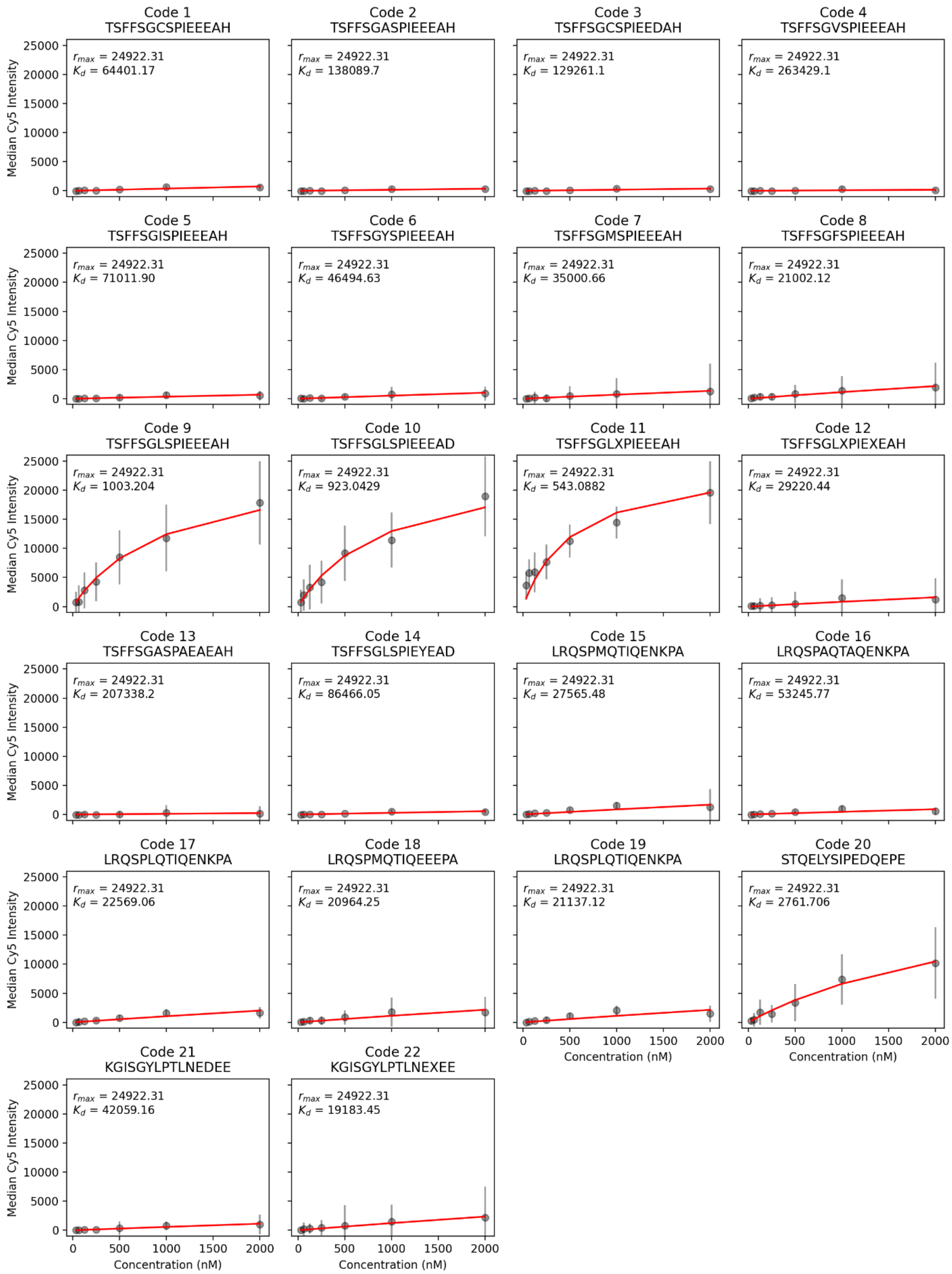


**Fig. S7**. Concentration-dependent binding data (grey markers, median intensity and standard deviation over all beads) and associated Langmuir isotherm fits (red lines) for MRBLE-pep library 1, replicate #2.

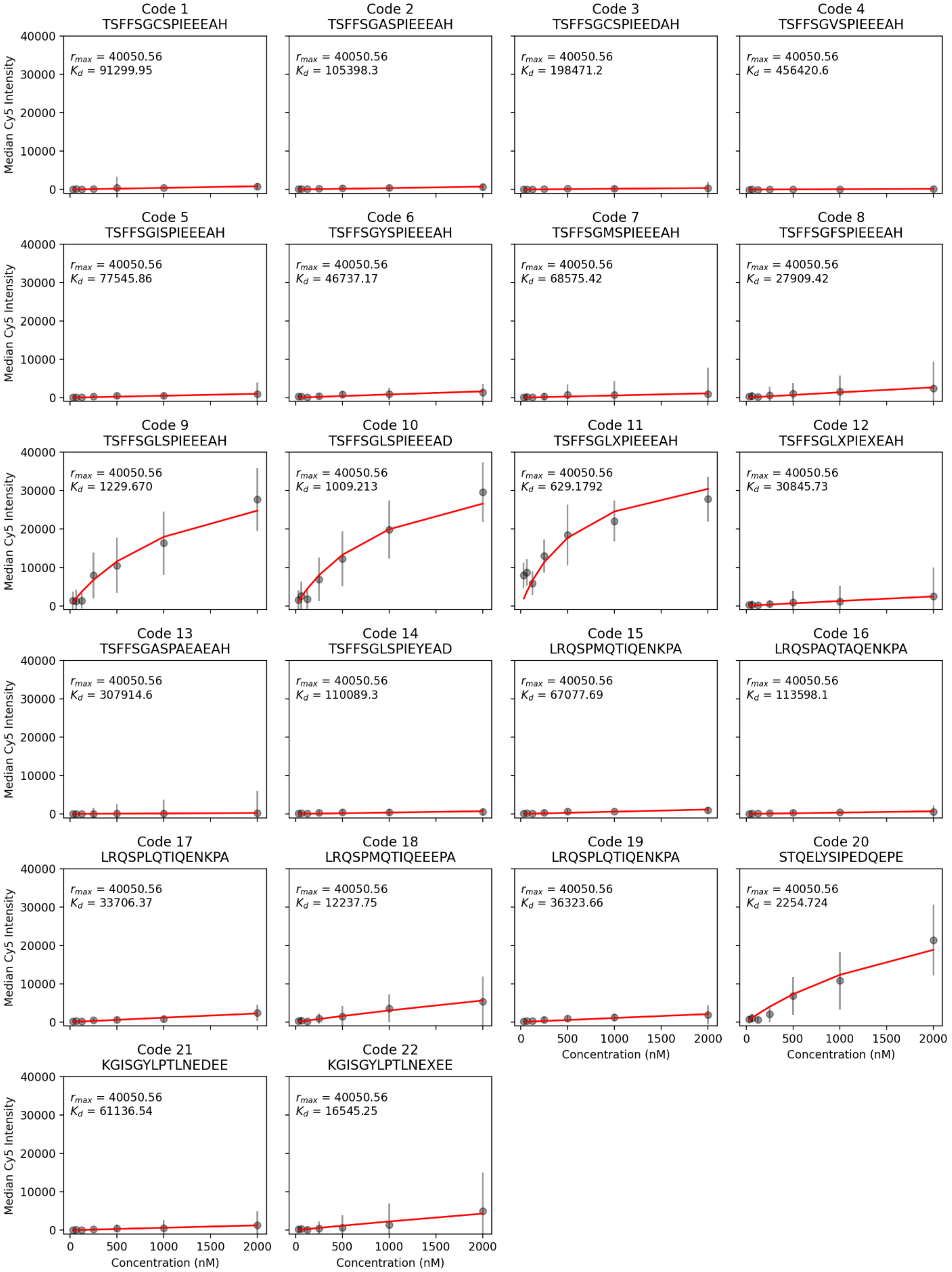


**Fig. S8**. Concentration-dependent binding data (grey markers, median intensity and standard deviation over all beads) and associated Langmuir isotherm fits (red lines) for MRBLE-pep library 1, replicate #3.

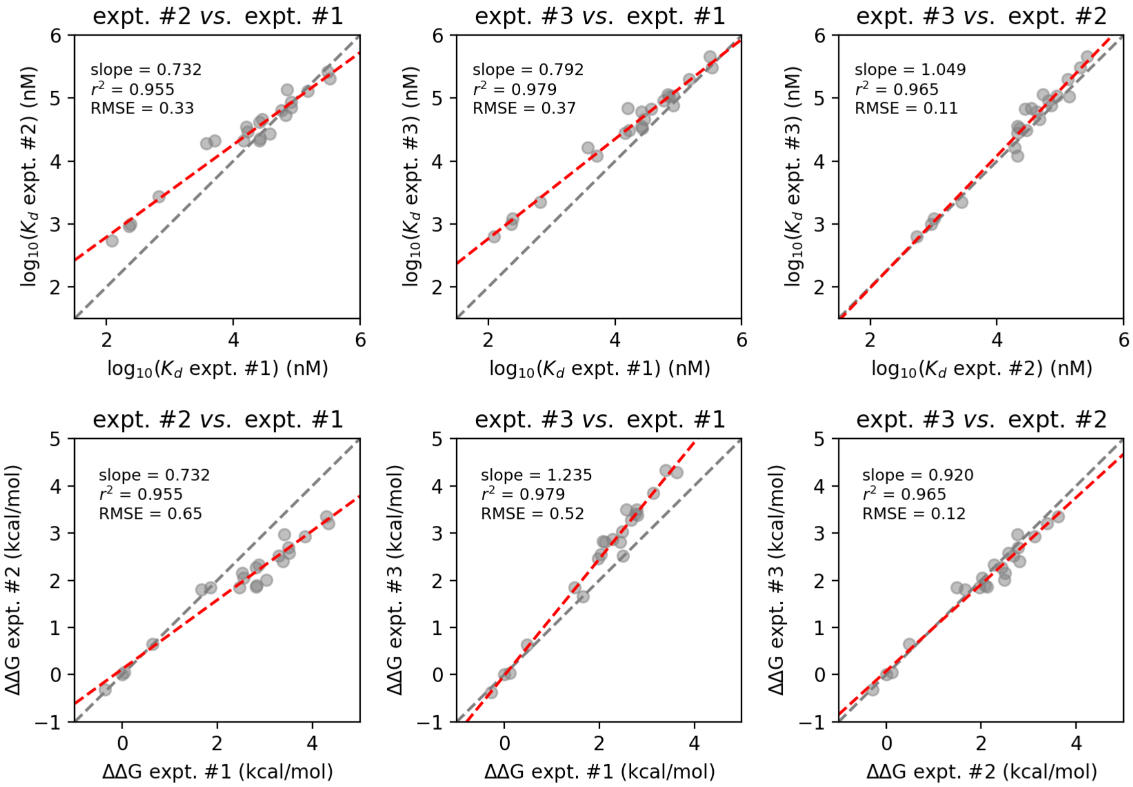


**Fig. S9**. Comparison of measured affinities between technical replicates for MRBLE-pep library 1. **(A)** Comparison of $K_d$s for each B56-peptide interaction (returned values from global Langmuir isotherm fits). Black dashed line indicates the 1:1 identity line; red dashed line indicates a linear regression to $\log_{10}$-transformed $K_d$ values. (**B**) Comparison of ΔΔGs for each B56-peptide interactions (calculated relative to a Kif4A 'reference' peptide sequence of TSFFSGLSPIEEEAD). Black dashed line indicates the 1:1 identity line; red dashed line indicates a linear regression.

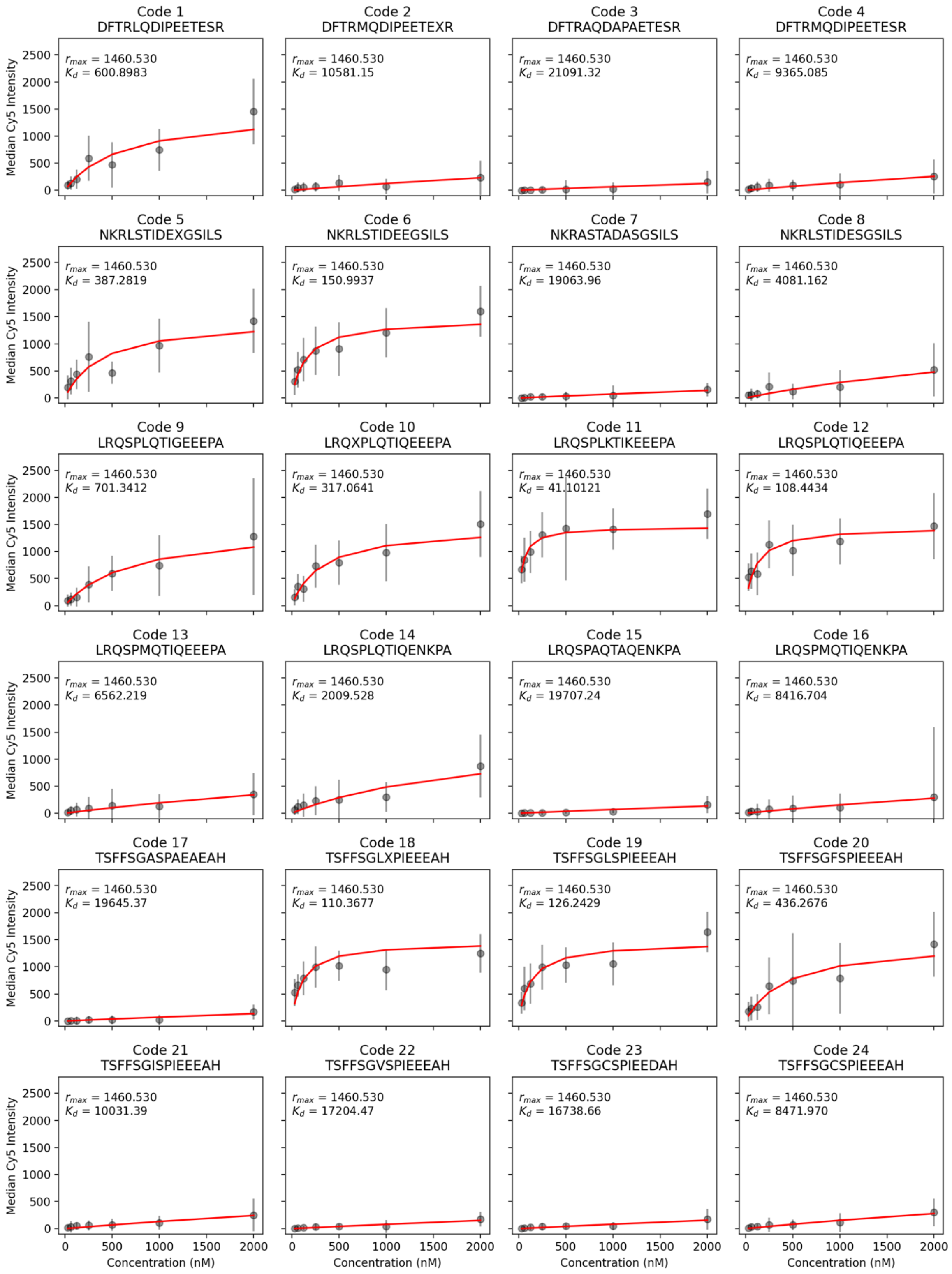


**Fig. S10**. Concentration-dependent binding data (grey markers, median intensity and standard deviation over all beads) and associated Langmuir isotherm fits (red lines) for MRBLE-pep library 2, replicate #1.

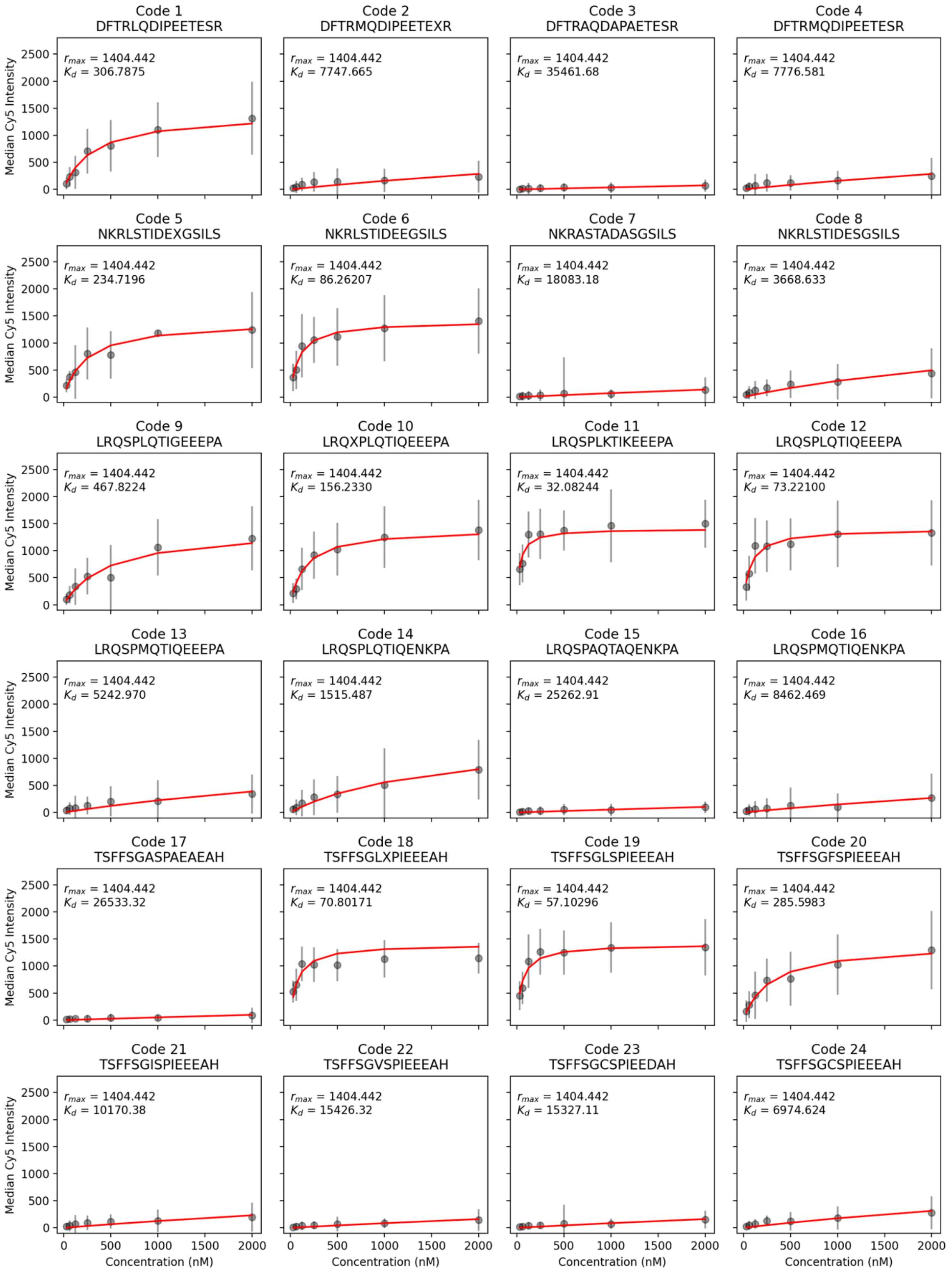


**Fig. S11**. Concentration-dependent binding data (grey markers, median intensity and standard deviation over all beads) and associated Langmuir isotherm fits (red lines) for MRBLE-pep library 2, replicate #2.

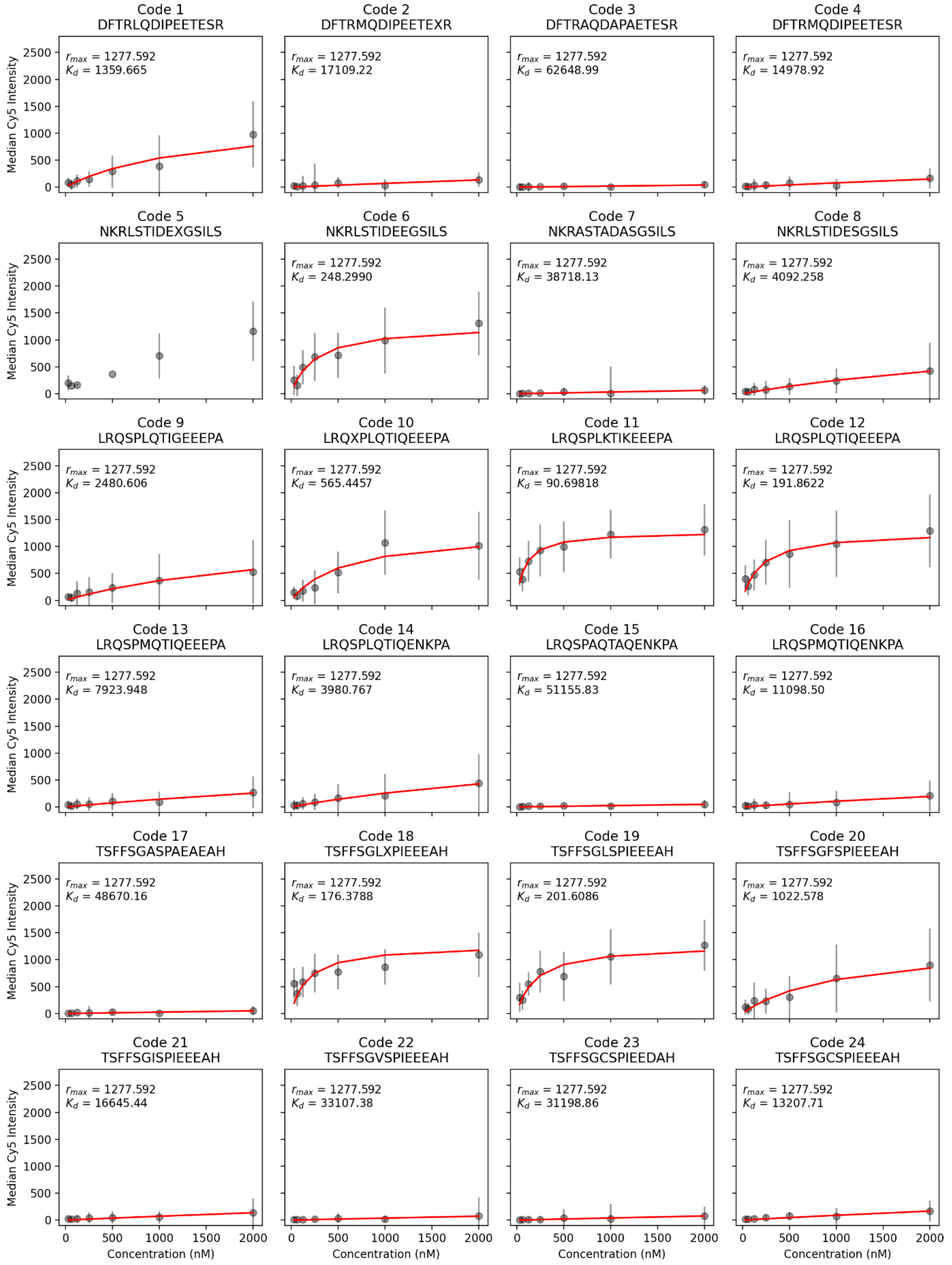


**Fig. S12**. Concentration-dependent binding data (grey markers, median intensity and standard deviation over all beads) and associated Langmuir isotherm fits (red lines) for MRBLE-pep library 1, replicate #3.

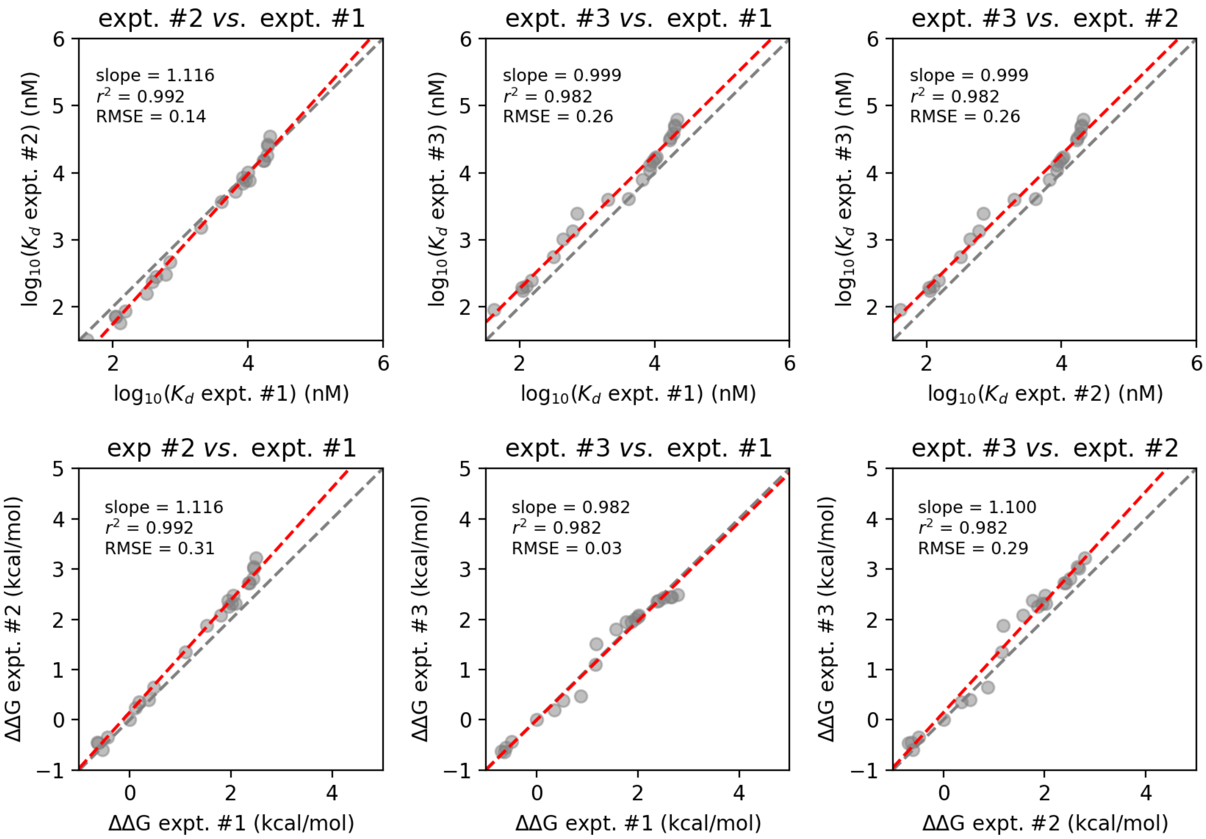


**Fig. S13**. Comparison of measured affinities between technical replicates for MRBLE-pep library 2. **(A)** Comparison of $K_d$s for each B56-peptide interaction (returned values from global Langmuir isotherm fits). Black dashed line indicates the 1:1 identity line; red dashed line indicates a linear regression to $\log_{10}$-transformed $K_d$ values. (**B**) Comparison of ΔΔGs for each B56-peptide interactions (calculated relative to a Kif4A 'reference' peptide sequence of TSFFSGLSPIEEEAD). Black dashed line indicates the 1:1 identity line; red dashed line indicates a linear regression.

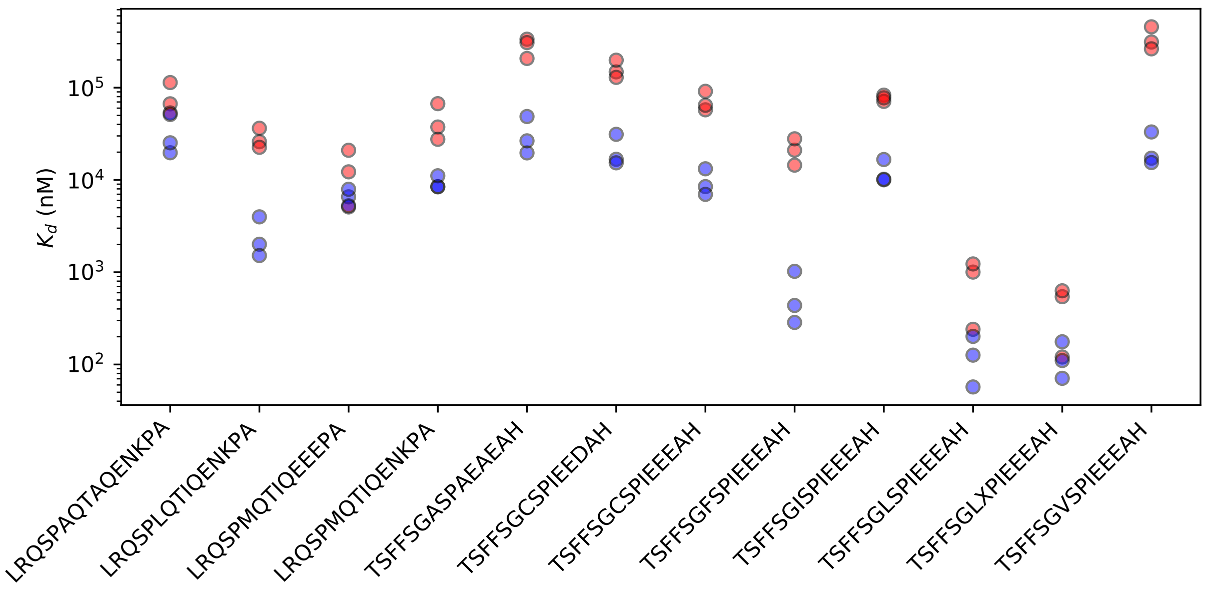


**Fig. S14**. Measured $K_d$s for 12 peptides containing Kif4A-like or FoxO3-like motifs across 3 technical replicates each of MRBLE-pep library 1 (red markers) and library 2 (blue markers).
